# A hydrogel beads based platform for single-cell phenotypic analysis and digital molecular detection

**DOI:** 10.1101/848168

**Authors:** Yanzhe Zhu, Jing Li, Xingyu Lin, Xiao Huang, Michael R. Hoffmann

## Abstract

Microfluidic platforms integrating phenotyping and genotyping approaches have the potential to advance the understanding of single cell genotype-to-phenotype correlations. These correlations can play a key role in tackling antibiotic heteroresistance, cancer cell heterogeneity, and other related fundamental problems. Herein, we report a novel platform that enables both high-throughput digital molecular detection and single-cell phenotypic analysis, utilizing nanoliter-sized biocompatible polyethylene glycol hydrogel beads produced by a convenient and disposable centrifugal droplet generation device. The hydrogel beads have been demonstrated enhanced thermal stability, and achieved uncompromised efficiencies in digital polymerase chain reaction, digital loop-mediated isothermal amplification, and single cell phenotyping. The crosslinked hydrogel network highlights the prospective linkage of various subsequent molecular analyses to address the genotypic differences between cellular subpopulations exhibiting distinct phenotypes. Our platform shows great potential for applications in clinical practice and medical research, and promises new perspectives in mechanism elucidation of environment-evolution interaction and other basic research areas.

## Introduction

Microfluidic single cell techniques have enabled observations of rare genotypes or phenotypes within a cell population and thus ubiquitous cell heterogeneity (1-3). The phenotypic diversity exhibited by supposedly genetically identical cells boosts the population adaptability under selection pressures, and thus raises concerns in fields spanning from clinical practice to medical research on infectious diseases and cancers (4, 5), etc. For example, less susceptible pathogenic bacterial subpopulations originally consist 10^-2^ to 10^-6^ of the overall population that can be amplified during antibiotic exposure. The subsequent increase in the resistant subpopulation may eventually lead to the failure of an antibiotic treatment (6). Hypotheses for the underlying molecular mechanisms involving the stochasticity of genetic mutation, gene expression, and protein regulation (7-9), however, remain hard to test in dynamically changing cell subpopulations, partly due to the absence of appropriate single cell experimental technique (10). The need to better understand cell heterogeneity motivates the development of new techniques that link the single-cell phenotype with its *in situ* molecular information.

As an emerging class of technologies, water-in-oil droplet-based microfluidic platforms have been well developed for high-throughput phenotypic and molecular analyses at single cell or single molecule resolution (3, 11). Nonetheless, due to the rare and transient nature of cell heterogeneity events, population-averaged molecular analyses would most likely fail to directly explain the characterized phenotypes, even if all analyses are conducted at single cell or molecular resolution (6, 12). Meanwhile, incorporating a crosslinked hydrogel network into the aqueous phase theoretically provides a droplet-based platform with additional robustness by allowing reagent exchange (13). This strategy, therefore, has been explored for a range of hydrogel materials and crosslinking chemistry, including cooling-induced formation of agarose beads for digital droplet polymerase chain reaction polymerase chain reaction (ddPCR) (14), ionic crosslinking of alginate beads for cell encapsulation and DNA extraction (15, 16), UV-initiated polyethylene glycol (PEG) beads for cell encapsulation (17). Such platforms have demonstrated to be effective in either phenotyping or molecular analysis, while the material and/or initiation method would be intrinsically incompatible with the combination of both. For example, temperature manipulation or UV radiation might affect the phenotype and genotype of encapsulated cells (18), and alginate is a well-known PCR inhibitor (19). PEG crosslinked by a thiol-Michael addition reaction between the bioinert acrylate and thiol groups has been attempted in bulk analyses and is among the most promising solutions (20, 21), but it is yet to be developed for our specific purpose. The main obstacle may lie in the fast and spontaneous gelation, which would be detrimental to traditional expensive microfluidic droplet generation approaches.

Herein, we report a novel PEG hydrogel bead-based platform, which is validated for both single-cell phenotypic analysis and molecular detection (**Figure 1a**-**b**). To solve the challenge posted by the fast thiol-Michael addition gelation chemistry, we developed a disposable centrifugal device for droplet generation (**Figure 1c**). We demonstrated the effectiveness of nucleic acid amplification detections, including PCR and loop-mediated isothermal amplification (LAMP), through further crosslinking generated droplets into PEG hydrogel as PEG hydrogel beads (Gelbeads). Compared to ddPCR and ddLAMP, Gelbead-based digital PCR and LAMP (gdPCR and gdLAMP) were found to exhibit enhanced thermal stabilities and uncompromised amplification efficiencies. Gelbeads were also demonstrated effective for single cell encapsulation and phenotyping within 4 hr for tested bacteria. We envision that this platform will be of broad interest to researchers from many fundamental fields. The Gelbead platform reported here for the first time promises unprecedented capabilities for investigation of cell heterogeneity.

**Figure 1.**
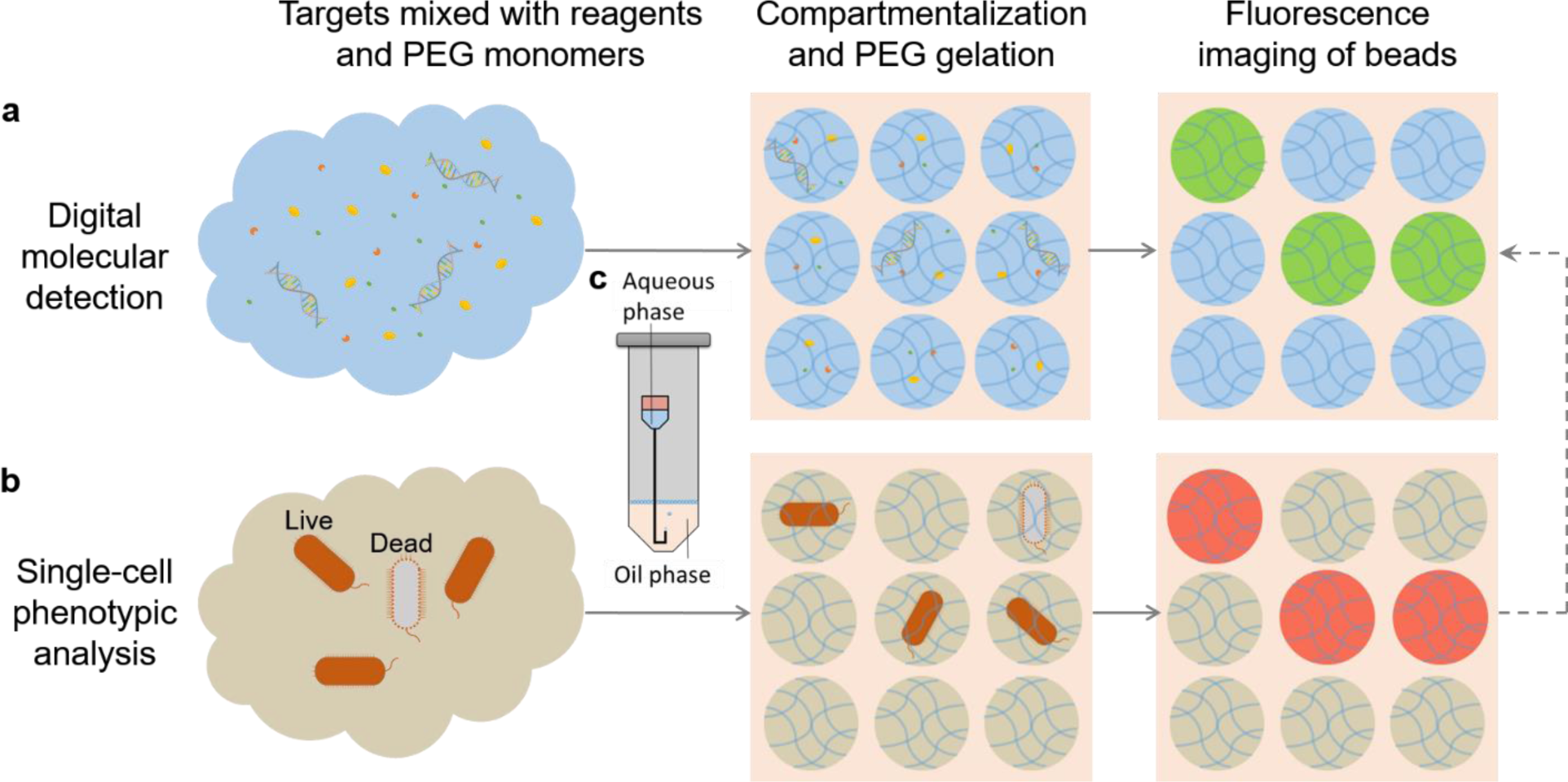
Schematic of this study. A hydrogel bead (Gelbeads)-based cell analysis platform was developed for (a) digital molecular detection including PCR and LAMP and (b) single-cell phenotypic analysis. The compartmentalization was realized by (c) a disposable centrifugal droplet generation device. The dashed-line arrow indicates that the crosslinked hydrogel network grants the potential of linking cell phenotype with *in situ* DNA/RNA characterization at single-cell resolution.

## Results

### Development of the disposable droplet generation device

Microfluidic-based droplet generation methods generally require special fabrication facilities to fabricate sub-100 µm channels and involve complicated operation, such as syringe pump-driven T-junctions fabricated by photolithography and centrifugally driven labs-on-a-disc fabricated by micro milling and hot embossing (22, 23). These traditional methods are not compatible with Gelbead generation due to fast clogging imposed by the thiol-Michael addition chemistry. The bulk PEG crosslinking experiments showed that the time frame for droplet generation before gelation was as short as 8.5 min with the chosen hydrogel concentration at 7.5 w/v% (**Supplementary Note 1**, **Table S1**). In order to easily generate Gelbeads within minutes without clogging expensive microfluidic equipment, we designed a disposable device using affordable commercial components (**Figure 2a**). The device utilized a dispensing blunt needle with a bent tip. The bent-tipped needle was then set into a 1.5-mL microcentrifuge tube with oil to establish the physics for centrifugal droplet generation. With centrifugal acceleration, the aqueous phase is forced into the fluorinated oil phase by the elevated pressure difference between the reservoir surface and the narrow inlet. The fluorinated oil phase with a higher density pinches off the aqueous droplets, which then float to the air-oil interface.

**Figure 2.**
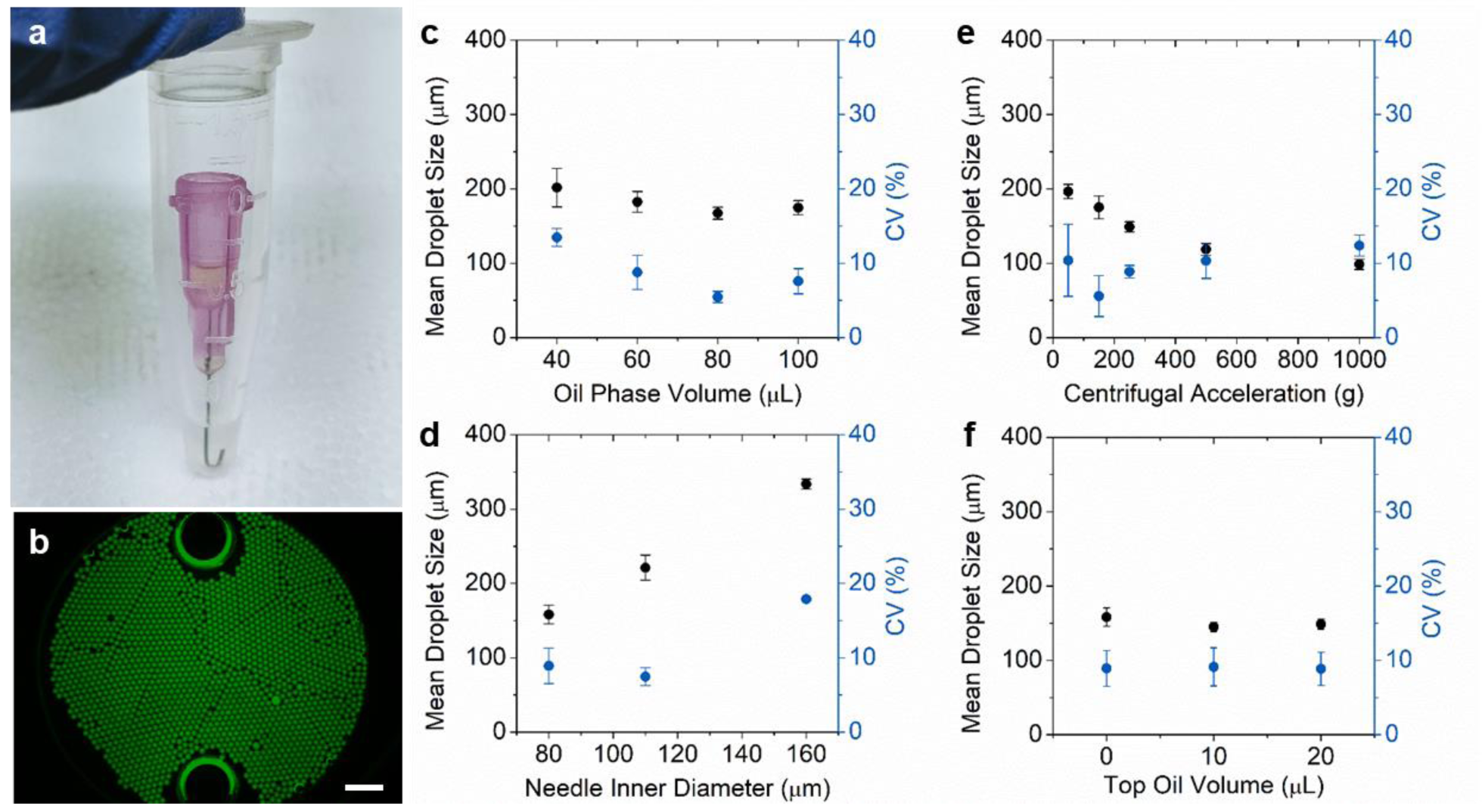
Development and evaluation of disposable microfluidics for centrifugal droplet generation. (a) The device setup consisting of a 1.5-mL microcentrifuge tube holding the oil phase and a needle with bent tip holding the aqueous reaction mixture in the Luer-lock. (b) A representative fluorescence microscope image of generated droplets extracted into a viewing chamber. The two large bright circles are ports on the viewing chamber for liquid loading Scale bar, 1 mm. (c-f) Mean droplet size (black circles) and CV (blue circles) of droplets produced under varying parameters including (c) oil phase volume, (d) needle inner diameter, (e) centrifugal acceleration and (f) oil volume added to the Luer-lock. Error bars represent standard deviation from independent triplicates.

Standard 20 µL LAMP mix with unquenched calcein was dispersed in fluorinated oil (online methods) and characterized using a fluorescence microscope to study the droplet generation performance of the device (**Figure 2b**). The average droplet size was tunable from 99 µm to 334 µm and the coefficient of variance (CV) was minimized to 5%, by varying the oil phase volume, centrifugal acceleration, and the needle gauge as shown in **Figure 2c**-**f**. Smaller droplets with slightly larger size distribution (**Figure 2e**) were produced by increasing the centrifugal acceleration, which provided a greater pressure difference to drive the aqueous phase inflow. The larger CV in **Figure 2e** was likely due to the unstable flow during initial acceleration, which can be alleviated by adding more oil (**Figure 2c**) to reduce the oil phase height variation and limit the amount of aqueous phase inlet during acceleration. Among all tested conditions, the optimal was found to be a combination of 34 Ga needles, 80 µL oil phase, and 150 g centrifugation run for 5 min and droplets were produced at an average diameter of 175 µm in 5 min with minor trial-to-trial difference, which was found to be comparable to other microfluidic methods such as centrifugal lab-on-a-disk (22) and polymer-tube micronozzles (24) (**Supplementary Note 2**). For droplets of a diameter of 175 µm, each standard 20 µL reaction could theoretically produce ∼10^4^ droplets. Based on this calculated compartmentalization, the dynamic range is theoretically from 0.5 to 3×10^3^ target copies or cells per µL, and the detection limit is 0.1 copies or cells per µL (25).

### Gelbead generation and thermal stability characterization

The Gelbead and droplet generation performance were assessed using various reaction matrices including culture media, PCR mix, and LAMP mix, under the optimized condition reported in the previous section (**Figure 3a**). The average diameter of generated Gelbeads was found to range from 145 µm to 217 µm with a CV from 3.6 % to 7.6 %. The observed variations were likely due to viscosity differences and interfacial property changes in different reaction matrices. It should be noted that the culture media alone was not able to sustain as droplets or Gelbeads in the fluorinated oil by 5% FluoroSurfactant. Bovine serum albumin (BSA), a protein commonly used as an additive to protect essential molecules (fatty acids, amino acids, etc.) in culture media (26), was added to the aqueous phase as an additional surfactant to modify interfacial properties and thus prevent the droplet merging. For the PCR reaction matrix, the generated Gelbeads had a larger CV than droplets. We assume that the presence of PEG hydrogel may have disturbed the surfactant-stabilized aqueous-oil interface, by inducing interfacial adsorption of additional charged species such as thiolate, magnesium ions, etc. In summary, the observed sizes and CVs of droplets and Gelbeads were considered acceptable for our assays. In general, this generation device fulfills the requirements for Gelbead generation. The simple generation device may be used for applications for which a simple yet powerful compartmentalization method is needed.

**Figure 3.**
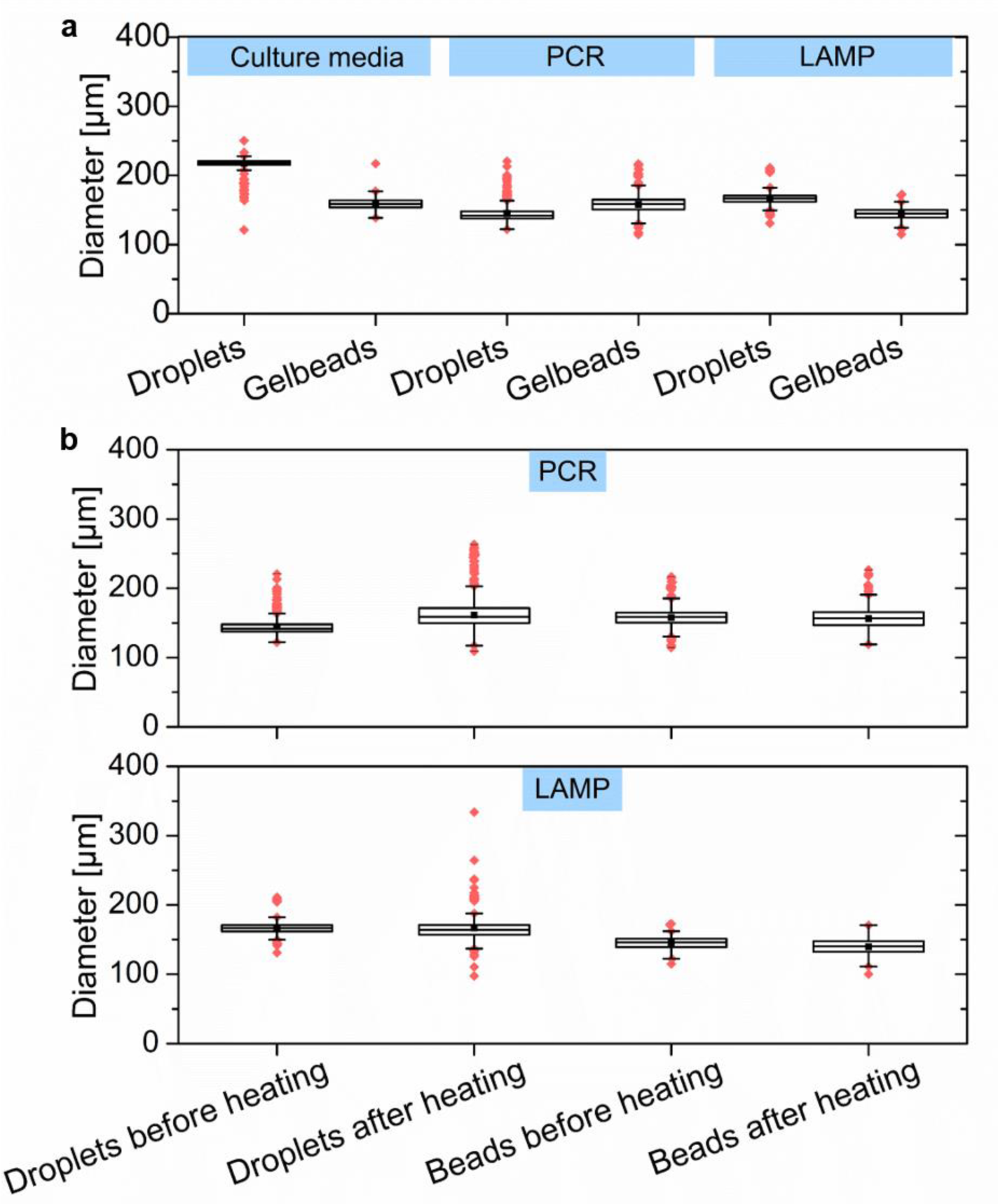
Size characterization of droplets and Gelbeads. The size distribution of droplets and Gelbeads (a) generated in reaction matrices including PCR mix, LAMP mix, and culture media mix, and (b) before and after heating program designated for PCR and LAMP. The line inside each box represents the mean diameter; the lower and upper edges of each box respectively represent 25% and 75% percentiles; the vertical bars below and above each box respectively indicate 90^th^ and 10^th^ percentiles. The lower and upper red dots stand for outliers.

The effect of PEG crosslinking on stabilizing the aqueous-in-oil compartments was evaluated. Thermodynamic instability of water-in oil droplets may impair the reliability of amplification processes such as PCR and LAMP that require extensive heating (22). Heating accelerates droplet merging and evaporation, which would affect the fluorescence reading by modifying concentrations of targets and reagents (e.g., salts and fluorescent dyes). The size distributions were investigated for droplets and Gelbeads before and after common heating protocols respectively for PCR and LAMP (online methods, **Figure 3b**). Compared to those before heating, droplets that had undergone PCR and LAMP heating increased in their CVs by 6.2% and 3.5%, respectively. In addition, the heating resulted in a noticeably larger population with outlier sizes implying that extensive merging and evaporation had occurred. Following the same heating protocol as for the droplets, the Gelbeads exhibited much less of a change in size distribution (CV increased by 1.9% for PCR and 1.6% for LAMP), however the average Gelbead diameter decreased slightly. These results indicate that the stabilization effect achieved by crosslinked PEG was mainly by prevention of the merging of beads. Gelbeads used for the LAMP procedure had a more significant improvement in thermal stability due to PEG crosslinking than for the PCR procedure. We assume that, in the case of the PCR recipe, the combination of SuperMix and the oil phase from BioRad were chemically well-optimized for interfacial stability, leaving limited room for improvement. This result therefore indicates that, other than modifying the surfactant composition or increasing surfactant concentration, hydrogel crosslinking could be an alternative strategy for maintaining the emulsion. Our results demonstrate that Gelbeads are a reliable platform for standalone heated digital analysis in terms of enhanced individual compartment integrity.

### Gelbead digital PCR (gdPCR)

To validate the reliability of gdPCR, we compared gdPCR to digital PCR performed in droplets generated from a commercial recipe (represented as ddPCR, hereinafter) for amplification efficiency with DNA extracted from cultured *Salmonella* Typhi (*S.* Typhi). Previous use of hydrogels and PCR utilized polyacrylamide in the form of either a bulk phase hydrogel membrane as a quasi-digital PCR platform (27) or using hydrogel beads as a substrate for surface coating of primers (28, 29), which is an approach opposite to our concept. To the best of our knowledge, performing PCR inside crosslinked hydrogel beads has not been reported to date. Even in bulk membrane form, only 80% amplification efficiency was observed, which may be partially attributed to template damage by free radicals as suggested (27). In this study, a similar drop in amplification efficiency was observed in the Gelbeads compared to that in droplets (**Figure 4a**), even though the Michael addition chemistry between acrylate and thiol used in this study does not involve free radical formation. In this case, crosslinked hydrogel network may be responsible for the observed inhibition by limiting the diffusion of functional components such as ions, nucleic acids, and proteins, where the extent of the limitation relates to the size and charge of the component (30, 31). From effective diffusivity modeling (**Figure S1**), we reasoned that the most affected functional component might be DNA polymerase, which is the relatively large protein (∼6 nm) responsible for building amplicons. For a fixed template concentration of 200 copies/µL estimated by ddPCR, gdPCR assay performance was assessed with additional OneTaq polymerase supplied at varying concentrations of 0.025, 0.05, 0.1, 0.2 Units per reaction, as shown in **Figure 4a**. Results showed that additional 0.025 Unit per reaction, 5% of the recommended OneTaq polymerase concentration per reaction, boosted the amplification efficiency the most. OneTaq polymerase concentrations supplied more or less than that showed inhibition to amplification efficiency, and gdPCR assay with additional 0.2 Unit per reaction was shown to be completely inhibited. We speculate that the observed trend was mainly due to the commercial SuperMix buffer conditions not optimized for the supplied OneTaq polymerase. While some additional polymerase compensated the reduced diffusivity of the SuperMix polymerase in hydrogel, the excess additional OneTaq polymerase might scavenge the essential ions for the original polymerase from SuperMix leading to amplification failure. With the optimized additional polymerase, gdPCR assays for serially diluted DNA with concentrations ranging from 2.5 to 600 copies/µL were then performed; typical images are shown in **Figure 4c**-h (**Supplementary Note 3**). The image analysis results demonstrated that the amplification efficiency of gdPCR was comparable (*k* = 0.98 ± 0.02, *R*^2^ = 0.9979) to that of ddPCR with the recipe adjustment (**Figure 4b**). The quantification results also correlated well with input DNA concentration (Figure S2a). It should be noted that the crosslinking inhibition effect eliminated in this case was for a 131 bp target gene (32), a typical size for detection of specific bacteria. Further optimization in polymerase or Supermix concentration would be required for other applications if a larger DNA fragment is targeted.

**Figure 4.**
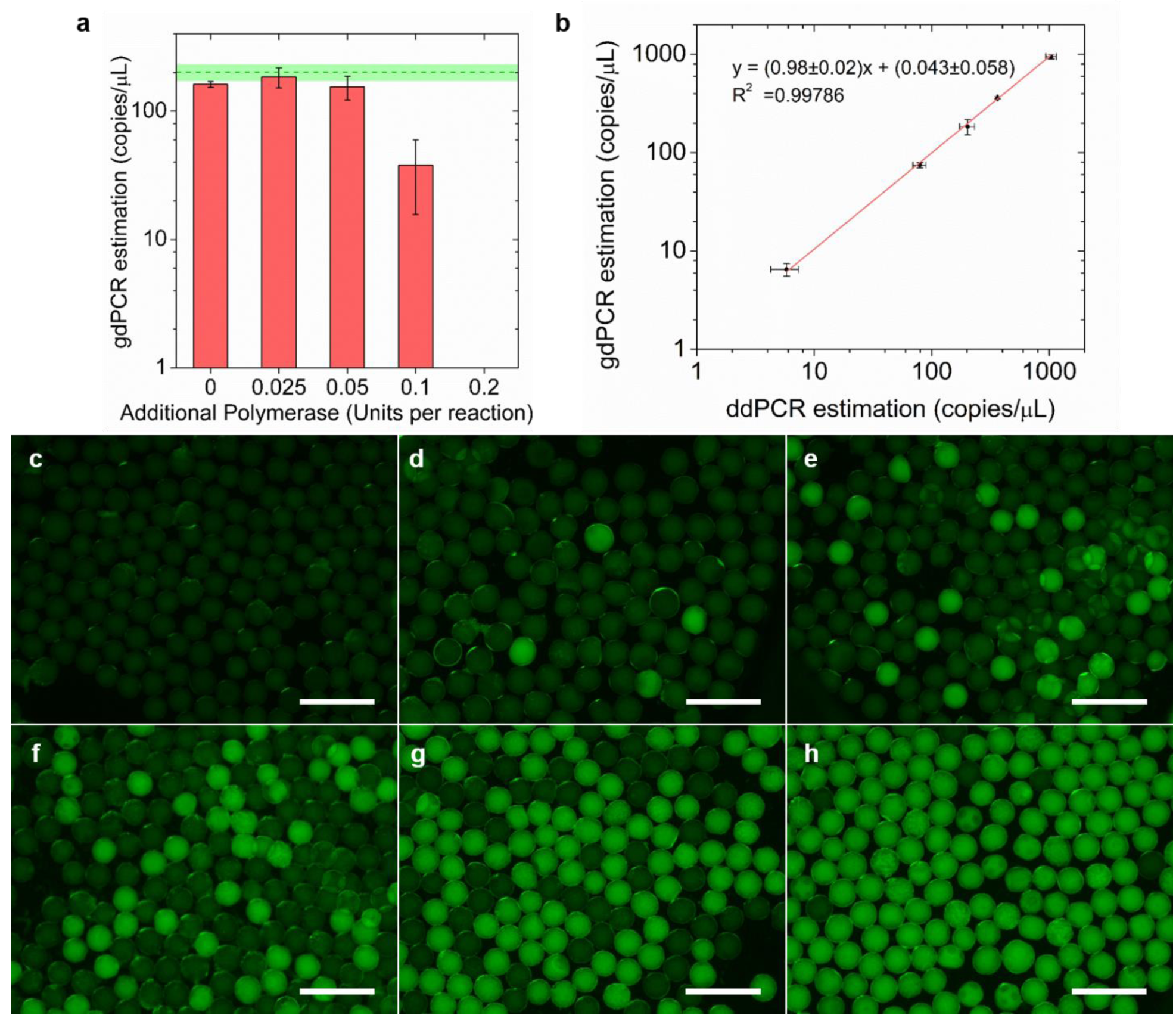
Optimization and performance of gdPCR. (a) The concentration estimations of gdPCR assays for a fixed input S. Typhi DNA concentration (200 copies/µL) with varying concentrations of additional polymerase. The green dashed line and the green area represent mean concentration estimation with standard deviation of ddPCR assays from independent triplicates. (b) With the optimized additional polymerase concentration (0.025 Units per reaction), the correlation between gdPCR and ddPCR estimation for serial diluted target templates. Error bars represent standard deviations from independent triplicates. (c-h) Example gdPCR fluorescent images for no DNA input, and with 24000, 1500, 600, 300, 100 times dilution of harvested S. Typhi DNA. Scale bars, 500 µm.

### Gelbeads digital LAMP (gdLAMP)

Gelbeads applied in digital LAMP were also investigated. LAMP has been an attractive emerging platform for molecular detection since it eliminates the need for thermocycling by utilizing a combination of 4 or 6 primers to achieve fast and specific detection (33). The heating protocol of LAMP was fairly mild, however, severe Gelbead aggregation occurred for samples with target DNA but not for no-template controls (**Figure S3**) in preliminary experiments. This was supposedly due to the fact that LAMP produces a much larger amount of amplification products than PCR (33).The negatively charged amplified DNA may have affected interfacial tension when adsorbed to the interface. Aggregated Gelbeads showed apparent crosstalking, which rendered the assay invalid since the compartment independence assumption required for Poisson statistics was contradicted. The problem was relieved by adding 1.5 mg/mL BSA, a common real-time PCR additive, to prevent surface adsorption. However, it was still observed that positive Gelbeads tended to stick next to each other (**Figure 5a**). The observed radiative patterns in Gelbeads manifested the differential diffusivity of amplification products of varying size in crosslinked hydrogel network. A similar radiative pattern was observed by Huang et al. in LAMP performed in a hydrogel membrane (21). In our case, neither of the two radiative centers were at the connected interface, indicating that the stickiness may not have led to false positive Gelbeads within the time frame tested. The connection of positive Gelbeads was most likely the result of a change in interfacial tension caused by large amount of the negatively charged DNA produced during amplification. Further crosslinking breaking through the oil barrier would only occur when the positive Gelbeads encounter each other. In summary, the connected interface should not affect the quantification results. The gdLAMP quantifications for no-template control and serial diluted *S.* Typhi DNA ranging from 300 to 1.2 × 10^4^ copies/µL were then verified. Example images are shown in **Figure 5c**-**h**. The image analysis results demonstrated that the amplification efficiency of gdLAMP was similar (*k* = 1.01 ± 0.01, *R*^2^ = 0.9996) to that of ddLAMP (**Figure 5b**). However, both ddLAMP and gdLAMP gave concentration estimations ∼2 orders of magnitude lower than input DNA concentration (**Figure S2b**). Further increases in the amplification efficiency would likely require an improved primer design, which is out of the scope of this study. In summary, the results confirmed our hypothesis that the stickiness of positive Gelbeads do not considerably affect gdLAMP quantification, and demonstrated that the hydrogel network had a negligible inhibition effect on the digital LAMP assays that were performed.

**Figure 5.**
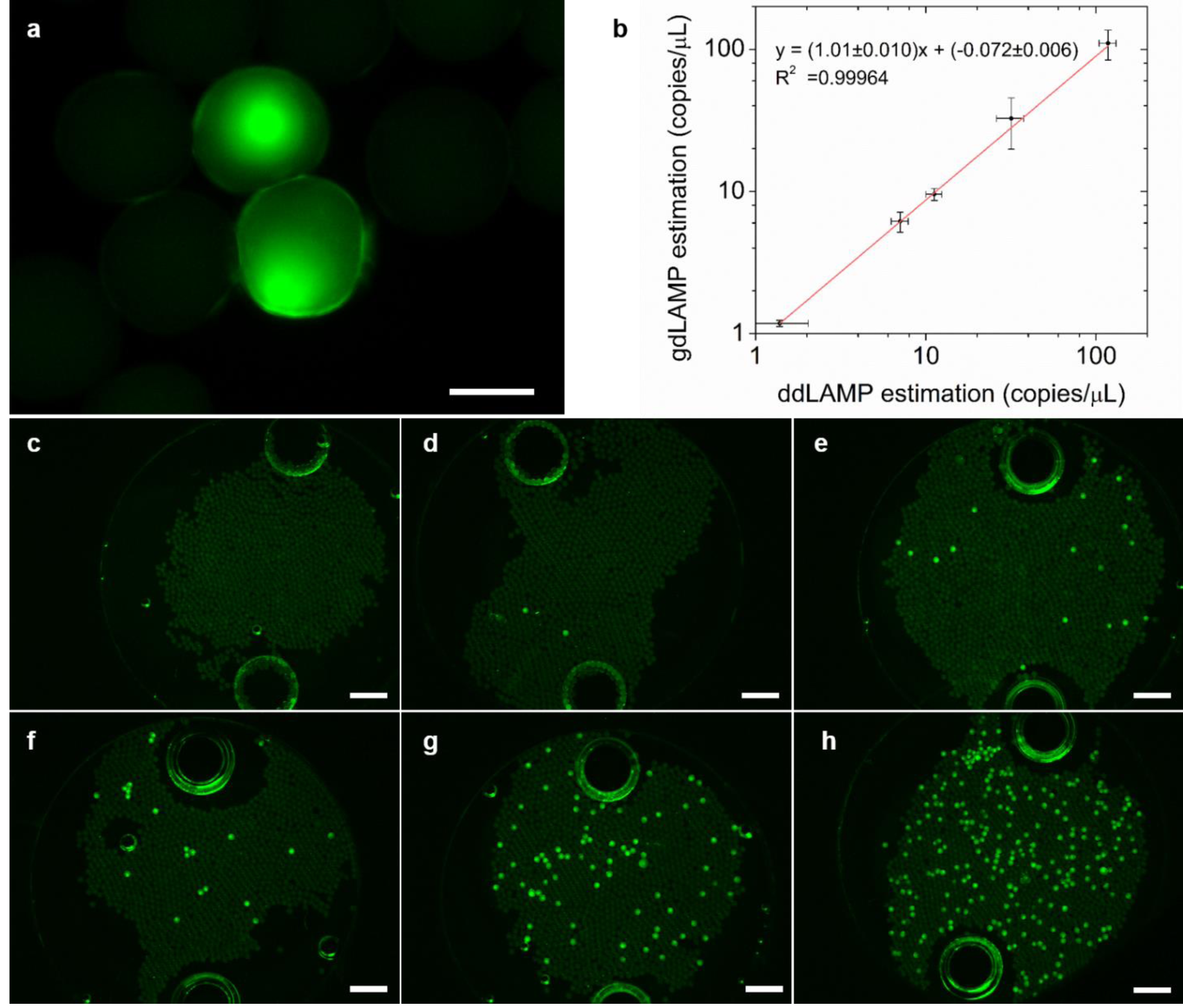
Performance of gdLAMP. (a) Connection of two positive Gelbeads after the gdLAMP assay. Scale bar, 100 µm. (b) The correlation between concentration estimations of gdLAMP and ddLAMP assays for serial diluted target templates. Error bars represent standard deviation from independent triplicates. (c-h) Example gdLAMP fluorescent images for no DNA input, and with 200, 100, 50, 20, 5 times dilution of harvested *S.* Typhi DNA. The two large bright circles on each image are ports on the viewing chamber for liquid loading. Scale bars, 500 µm.

### Gelbeads for cell phenotyping

For single cell phenotyping, we first validated single cell encapsulation efficiency using *Salmonella* Typhimurium with green fluorescent protein (GFP). The cells were diluted to an average of 1 cell per Gelbead for counting the number of cells in each Gelbead (**Figure 6b**). At this cell concentration, theoretically, 34% of the compartments were occupied by single cells, which was the maximum following a Poisson distribution, 29% of the compartments encapsulated more than 1 cell, and 37% of the compartments contained no cells. As shown in **Figure 6a**, the observed number of encapsulated cells was close to the theoretical distribution. Gelbeads with high cell numbers were slightly less than predicted, possibly because some cells were located out of focus when imaged in spherical compartments at a high microscope objective. Since high throughput detection of stained cells within spherical compartments droplets or Gelbeads was challenging for fluorescence microscope imaging, we chose to employ cell metabolism indicator dye in Gelbead phenotyping experiments. As a resazurin-based dye used in bulk phenotyping assays of a wide range of cell lines, alamarBlue can be reduced by actively metabolizing cells into resorufin, whose bright red fluorescence can stain the whole compartment for visualization (34). Phenotyping of *S.* Typhi in Gelbeads was investigated by co-incubation of alamarBlue and *S.* Typhi in the culture media. The fluorescence of Gelbeads was monitored during the incubation for up to 4 hrs (**Figure 6d-h**). It was observed that Gelbeads appeared to be much brighter than the droplets were before incubation (**Figure S4**); this was possibly due to additional reduction of resazurin by thiol group (35). We suppose that the interference by thiol groups would not affect the phenotyping results since the monomers were rigorously mixed and evenly distributed into Gelbeads. Gelbeads containing live cells would exhibit even brighter fluorescence in the presence of sufficient AlamarBlue. The quantitative performance of Gelbead phenotyping was verified by analysis of observed fractions of bright fluorescent Gelbeads (see online methods and **Figure S5** for thresholding) compared to the theoretical value, as shown in **Figure 6c**. 63% of Gelbeads were supposed to contain greater than or equal to 1 cell and thus to be bright. The observed positive fraction of 62.0 ±1.5% after 4 hours of incubation matched well with the theoretical value of 63%. It was also noticed that, after 3 hours of incubation, the positive Gelbead fraction was 36.4±8.1%, which corresponds well with the theoretical fraction of Gelbeads (29%) encapsulating more than 1 cell. Based on the linear response of alamarBlue to the number of cells within the compartment (36), our results reasonably indicate that effective single cell phenotyping in Gelbeads is achievable within 4 hrs. However, 5 hr incubation lead to overly bright fluorescence and 92.9±2.7% bright Gelbeads, which was likely attributed to excessive incubation and the diffusion of metabolized fluorescent resorufin across the aqueous-oil interfacial barrier. Our results indicate that the optimization of incubation time is a race between cross-talking and cell proliferation.

**Figure 6.**
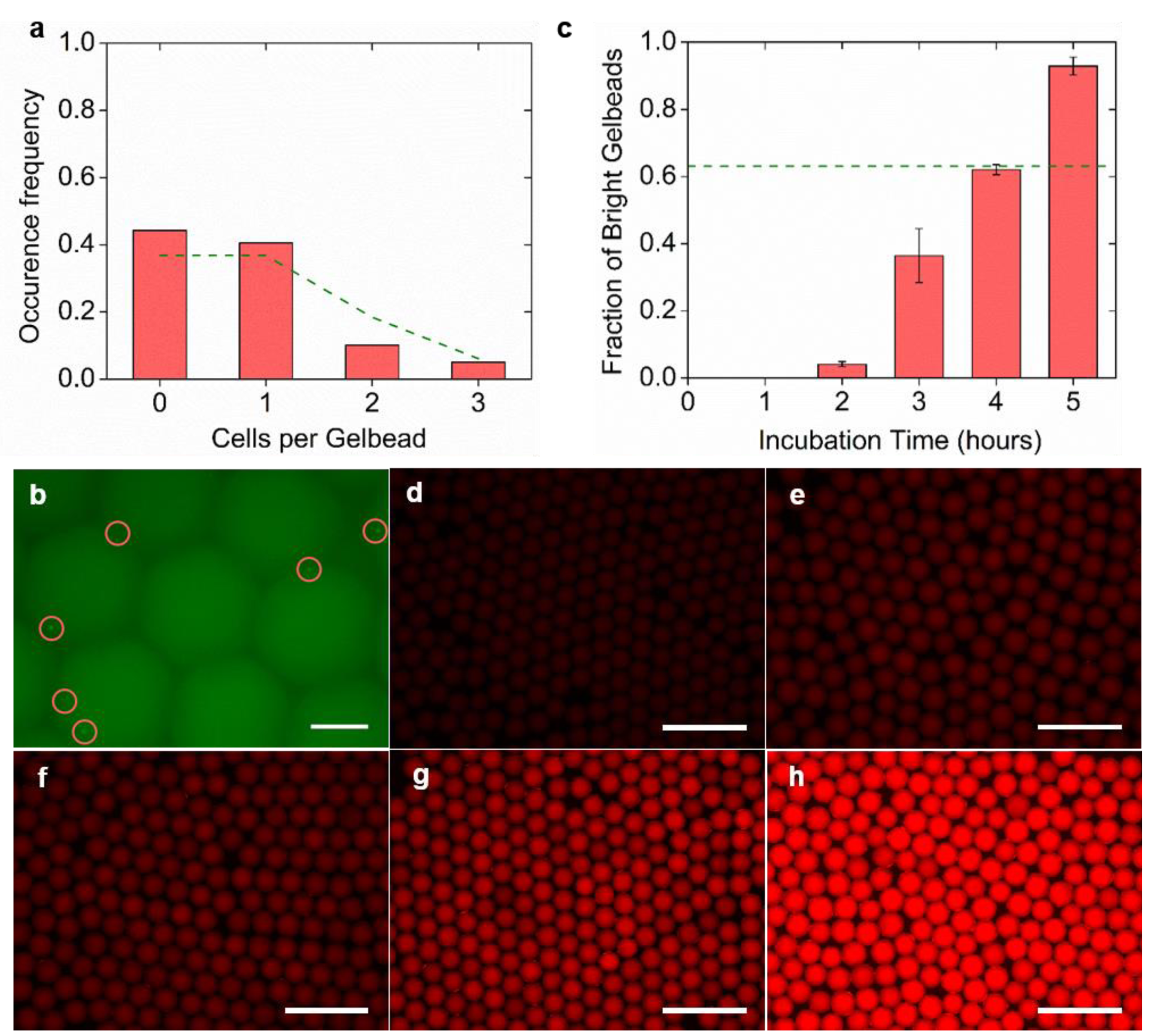
Single cell encapsulation validation and Single cell phenotyping performance in Gelbeads. (a) Number of cells encapsulated in each Gelbead counted and represented by occurrence frequency. The dashed line represent theoretical values based on Poisson distribution. (b) Example fluorescence image of encapsulated *S.* Typhimurium GFP cells (circled) for counting. Scale bar, 100 µm. (c) The observed fraction of bright Gelbeads with varying incubation time, with the dashed line representing 63% as Poisson distribution predicted based on the input cell concentration. Error bars represent standard deviation from independent triplicates. (d-h) Example images of Gelbeads containing *S*. typhi at the same input concentration incubated for 0, 2, 3, 4, 5 hrs. Scale bars, 500 µm.

Considering the intrinsic difference in proliferation rate between bacterial species, the observed incubation time for distinction of positive and negative compartments was comparable to the results by Lyu et al., who achieved *Escherichia coli* (*E. coli*) phenotyping with alamarBlue in 85 pL droplets with a 2 hr incubation (3). We note that, although the single cell encapsulated Gelbeads were maximized and theoretically comprised the majority (54%) of the bright Gelbeads in the current set up, strategies are available to break Poisson distribution for higher single cell encapsulation rates, such as microvortex-aided hydrodynamic trapping and then releasing single cells to droplets (37). In summary, the cell viability detection strategy demonstrated with Gelbeads has been proved to apply well to a wide range of cells in bulk assays and droplet microfluidics (3, 34, 36). Thus, the Gelbeads synthesized in this study provide a suitable platform for phenotyping cell heterogeneity, if they are co-encapsulated with antibiotics or drugs.

## Discussion

The developed Gelbeads platform promises a robust analysis tool that has the potential to link single-cell phenotypic analysis with reliable *in situ* molecular detection together. Besides the advantages presented, we acknowledge the following limitations. First, the dynamic range in our study was restricted by the size of the compartments generated by our device. Further reductions in size would result in larger size variations, and the surfactant might have to be changed or adjusted if higher uniformity is required. Second, given the use of fluorescence microscopic imaging of the compartments inside a viewing chamber, the Gelbead imaging approach employed could probe only a limited viewing area, and the resolution could be affected by the focus. The fluorescence characterization may be further improved by flow cytometry to interrogate single Gelbeads.

In this work, a disposable centrifugal device was developed for Gelbead generation using highly biocompatible PEG monomers spontaneously crosslinked with no free-radical, UV-induced or heat-induced initiation. Our design allows for easy use of droplet microfluidics without expensive and complicated equipment, which could useful for applications other than Gelbeads generation. In addition to the single cell phenotyping potential, the Gelbeads approach has enhanced thermal stability coupled with high amplification efficiency for dPCR and dLAMP. Widely available qPCR and LAMP assays can therefore be easily transferred into digital assays by this Gelbeads approach. The unique structural stability of the hydrogel network allows for easy manipulation of the Gelbeads that may have many possibilities for other upstream and downstream analyses. The Gelbead platform will be further developed for reagent exchange, fluorescence-based Gelbead sorting, and downstream sequencing, etc. We envision that the potential of our Gelbeads platform in generating genetic and gene expression data with phenotyped single cells will help narrow the genotype-phenotype gap and thus offer exciting new insights in cell heterogeneity studies.

## Materials and Methods

### PEG crosslinking and characterizations

PEG hydrogel monomers included 4-arm PEG-acrylate [molecular weight (MW) of 10 000, Laysan Bio, Arab, AL, USA] and thiol-PEG-thiol (MW of 3400; Laysan Bio), with acrylate and thiol mixed at a molar ratio of 1:1 for crosslinking. For sol-gel transition time characterization, 7.5 w/v% and 10 w/v% PEG hydrogel were respectively tested in PCR mix, LAMP mix, and culture media mix. PEG monomers were weighed to make 10× monomer solutions for PEG-acrylate and PEG-thiol separately. The weighed monomers were then dissolved either in water (Molecular Biology Grade Water, Corning, Acton, MA, USA) for PCR and LAMP mix, or in TSB (BD™ Bacto™ Tryptic Soy Broth, Becton Dickinson and Company, Franklin Lakes, NJ, USA) for culture media mix. In addition to 2 µL of each 10× PEG monomer solution, for each 20 µL reaction mix, PCR mix contained 10 µL ddPCR Supermix for Probes (BioRad, Hercules, CA, USA) and 6 µL water; LAMP mix contained 10 µL 2×WarmStart LAMP Mastermix (New England Biolabs, Ipswich, MA, USA) and 6 µL water; culture media mix contained 16 µL TSB. The reaction mix was briefly vortexed. The sol-gel transition was considered started when lifting the pipette tip could draw filaments out of the reaction mix, and the transition was considered ended when the reaction mix formed a gelatinous lump.

### Development of the disposable droplet generation device

Each droplet generation device consisted of a 1.5 mL DNA LoBind tube (Eppendorf, Hamburg, Germany) and a blunt tip dispensing needle (LAOMA Amazon, Seattle, WA, USA) with the tip bent by a tweezer (VWR, Radnor, PA, USA). The tweezer and the needles were autoclaved (2540EP, Heidolph Brinkmann, Schwabach, Germany) prior to use. The oil phase was added to the bottom of the microcentrifuge tube, and the aqueous reaction mix was added to the Luer-lock of the needle. The device was then centrifuged (Centrifuge 5430R, Eppendorf) for 5 min. For optimization of droplet generation, fluorinated oil (HFE-7500 3M^®^ Novec^®^ Engineering Fluid, 3M, Maplewood, MN, USA) supplied with 5% FluoroSurfactant (RAN Biotechnologies, Beverly, MA, USA) was added into the oil phase. The 20-µL aqueous phase contained 1×WarmStart LAMP Mastermix and 50 μM calcein (Sigma-Aldrich, St. Louis, MO, USA). Four parameters including oil phase volume, needle inner diameter, centrifugal acceleration and oil volume added to the Luer-lock were investigated. Specific variables in details were as follows: 1) the oil phase volume of 40, 60, 80 and 100 µL, respectively, at the bottom of the tube in 34 Ga needles under 250 g centrifugation; 2) needles of 30, 32 and 34 Ga (corresponding to inner diameter of around 160, 110 and 80 µm) under the condition of 250 g centrifugation and 80 µL oil phase volume; 3) the centrifugal accelerations of 50, 150, 250, 500, 1000 g with 34 Ga needles and 80 µL oil phase; 4) additional oil phase added into the Luer-lock of 0, 10 and 20 µL in 34 Ga needles under 250 g centrifugation with 80 µL oil phase.

### Gelbead generation and thermal stability characterization

In all the following experiments, the device configuration was fixed with 34 Ga needles, 80 µL oil phase, no additional oil at the Luer-lock, and 150 g centrifugation run for 5 min. The droplet and Gelbead generation using the described device was respectively characterized with PCR mix, LAMP mix, and culture media mix. In each 20 µL reaction, the PCR mix contained 1× ddPCR Supermix and 50 μM calcein; the LAMP mix contained 1×WarmStart LAMP Mastermix, and 50 μM calcein; the culture media mix was TSB with 1 mg/mL BSA (New England Biolabs) and 50 μM calcein. The mix was briefly pipette mixed. The reaction mix for Gelbead generation contained 7.5 w/v% PEG hydrogel, added as 10× PEG monomers. For dispersion of PCR mix as droplets and Gelbeads, Droplet Generation Oil for Probes (BioRad) was used instead of fluorinated oil with 5% FluoroSurfactant.

For thermal stability characterizations, generated droplets or Gelbeads were extracted into PCR tubes (0.2 mL individual PCR tubes, BioRad) and incubated in a thermal cycler (T100, BioRad). The thermocycling protocol for PCR included 10 min of initiation at 95 °C, followed by 40 cycles of denaturation at 94 °C for 30 s, annealing at 52 °C for 60 s, and extension at 65 °C for 30 s. For LAMP heating, droplets or Gelbeads were incubated at 65 °C for 1 hour.

### Bacterial Cell culture and DNA preparation

*Salmonella* Typhi (*S.* Typhi, CVD 909), obtained from American Type Culture Collection (ATCC, Manassas, VA, USA), was employed as the model strain. *S.* Typhi was cultivated in TSB supplied with 1 mg/L of 2,3-dihydroxybenzoate (DHB, Sigma-Aldrich) in an incubator (Innova 42, New Brunswick Scientific, Edison, NJ, USA) shaking at 200 rpm at 35 °C for 14-16 hours. The concentration of cultivated cells was estimated by OD 600 (NanoDrop 2000c Spectrophotometer, Thermo Scientific, Barrington, IL, USA). DNA was harvested using PureLink^®^ Genomic DNA Mini Kits (Fisher Scientific, Waltham, MA, USA) following the manufacturer’s instructions. For single cell encapsulation test, *Salmonella* Typhimurium GFP (ATCC 14028GFP) was cultivated in nutrient broth (Difco™ 23400, Becton Dickinson and Company) supplied with 100 mcg/ml Ampicillin (Sigma-Aldrich) in an incubator shaking at 200 rpm at 37 °C for 14-16 hours. The cell concentration was estimated by counting under a fluorescence microscope (Leica DMi8, Wetzlar, Germany).

### Gelbead Digital PCR (gdPCR) assay

The thermocycling protocol of gdPCR assay was the same as described in the thermal stability characterization. Each 20 µL reaction consisted of 1× ddPCR Supermix, 900 nM forward primer, 900 nM reverse primer, 250 nM probe, and 2 µL DNA sample or water. Additional 7.5 w/v% PEG hydrogel was added as 10× PEG monomers for gdPCR assays. The primers and probe were ordered from Integrated DNA Technologies (IDT, Coralville, IA, USA), with sequences (Supplementary Table S1) designed for specific detection of *S.* Typhi, targeting a region in gene STY0201 for an amplicon size of 131 bp (32). For gdPCR optimization, the same DNA template concentration (600 times dilution from harvested) was added for gdPCR assays and ddPCR control. Optimal concentration of additional polymerase (OneTaq^®^ DNA polymerase, New England Biolabs) was investigated by supplying various concentrations to the described reaction mix incrementally at 0.025, 0.5, 0.1 and 0.2 U/reaction. For quantification assays, harvested DNA sample were serial diluted 100, 300, 600, 1500, and 24000 times for ddPCR and gdPCR. The reactions were prepared on iceblock (Carolina^®^ Chill Block, Burlington, NC, USA) and centrifugation temperature was set at 4 °C. Droplets or Gelbeads were generated in BioRad droplet generation oil, and were then extracted into PCR tubes for thermocycling. No-template controls were examined for each tested condition.

### Gelbead Digital LAMP (gdLAMP) assay

The reagents for LAMP were acquired from New England BioLabs if not indicated otherwise. Each 20 μL of modified LAMP mix for digital single bacteria LAMP contained 1× isothermal buffer, 6 mM total MgSO4, 1.4 mM dNTP, 640 U/mL Bst 2.0 WarmStart polymerase, 1.6 μM FIB and BIP, 0.2 μM F3 and B3, 0.8 μM LF and LB, 1.5 mg/mL BSA, 1× LAMP dye (38). For gdLAMP assays, 7.5 w/v% PEG hydrogel was added as 10× PEG monomers. The primers, ordered from IDT with the sequences shown in Supplementary Table S1, were targeting a 196 bp region within the *S.* Typhi specific gene STY1607 (39). For gdLAMP and ddLAMP assays, harvested DNA was serial diluted 5, 20, 50, 100, and 200 times. The reactions were prepared on iceblock and centrifuged into 5% FluoroSurfactant supplied fluorinated oil at 4 °C. Droplets or Gelbeads were then extracted into PCR tubes for 30 min heating at 65 °C followed by 5 min polymerase deactivation at 80 °C. No-template controls were examined under the same protocol.

### Single cell phenotyping

For single cell encapsulation efficiency test, the cultivated *Salmonella* Typhimurium GFP was diluted 600 times for Gelbeads generation. The dilution factor was estimated from prior knowledge of cultured cell concentration and Gelbead volume. The number of cells encapsulated in each Gelbead was analyzed by fluorescence microscope imaging with a 20× objective. 79 Gelbeads were analyzed from 15 fluorescent images. For phenotyping experiments, 1mL of overnight cultured *S.* Typhi was freshly cultivated for 3 hours in 5mL TSB supplied with 1 mg/L of DHB in an incubator shaking at 200 rpm at 35 °C. The cell concentration was verified to be around 0.135 by OD 600. AlamarBlue (Invitrogen, Carlsbad, CA, USA) was employed as the cell viability indicator. To address the fluctuation of excitation intensity and emission detection within a microscopic view, calcein was used as a reference dye. Each 20 μL reaction consisted of 1× AlamarBlue, 50 μM calcein, 1 mg/mL BSA, diluted *S.* Typhi cells, and the rest of the volume filled with DHB supplied TSB. 7.5 w/v% PEG hydrogel was added as 10× PEG monomers dissolved in DHB supplied TSB. After generation, the Gelbeads were incubated at 37 °C for 0-5 hrs. Gelbeads were extracted for imaging after 0, 1, 2, 3, 4 hrs of incubation.

### Droplets and Gelbeads imaging and analysis

The droplets or Gelbeads to be analyzed were pipetted into a viewing chamber made by adhering SecureSeal™ Hybridization Chamber (9 mm DIA × 1.0 mm Depth, Grace Bio-Labs, Bend, OR, USA) to a glass slide (VistaVision^®^ Microscope slides, VWR). The chambers were imaged under the fluorescence microscope using a 1.25× objective for droplets/Gelbeads generation, characterizations, and gdLAMP. For each sample in gdPCR and single cell phenotyping, five images of different area in the viewing chamber were taken using a 5× objective. Fluorescein isothiocyanate (FITC) filter was used, except for phenotyping experiments where Texas Red (TXR) filter was used in addition. In phenotyping experiments, the image data collected through TXR channel was normalized using the image data collected through FITC channel. For analysis of bright Gelbeads fraction, the data of each pixel was the intensity ratio of TXR channel to FITC channel. All images were analyzed using customized MATLAB scripts (**Supplementary Files**). For droplets and Gelbeads generation as well as thermal stability characterizations, the images were analyzed for individual compartment diameters. The diameters were further analyzed to calculate average compartment diameter and coefficient of variation (CV). For gdPCR, gdLAMP, and phenotyping assays, in addition to size analysis, the images were also analyzed for number of positive and negative compartments by setting a bright-dark threshold. Using the ratio of negative compartments to total compartments, the input DNA or cell concentrations were estimated by Poisson distribution (40). For images from phenotyping assays, since the distinction of dark and bright Gelbeads was hard to inspect visually, Gaussian fitting was used to advice the threshold (**Figure S5**).

## General

We thank Dr. Katharina Urmann for helpful discussions.

## Author contributions

The manuscript was written through contributions of all authors. M.R.H, X.H., and Y.Z. conceived the concept for this study. J.L., X.H., X.L. and Y.Z. designed the study, Y.Z. performed experiments, and J.L. and Y.Z. wrote the paper. All authors approved of the manuscript.

## Data and materials availability

The manuscript and the supplementary materials contain all data needed to evaluate the conclusions in the paper. Correspondence and requests for materials should be addressed to M.R.H.

## Reference

1. C. M. O’Keefe et al., Facile profiling of molecular heterogeneity by microfluidic digital melt. Sci Adv 4, (2018).

2. L. F. Cheow et al., Single-cell multimodal profiling reveals cellular epigenetic heterogeneity. Nat Methods 13, 833–836 (2016).

3. F. J. Lyu et al., Phenotyping antibiotic resistance with single-cell resolution for the detection of heteroresistance. Sensor Actuat B-Chem 270, 396–404 (2018).

4. I. El Meouche, M. J. Dunlop, Heterogeneity in efflux pump expression predisposes antibiotic-resistant cells to mutation. Science 362, 686-+ (2018).

5. D. Shibata, Cancer. Heterogeneity and tumor history. Science 336, 304–305 (2012).

6. D. I. Andersson, H. Nicoloff, K. Hjort, Mechanisms and clinical relevance of bacterial heteroresistance. Nat Rev Microbiol, (2019).

7. U. Ben-David et al., Genetic and transcriptional evolution alters cancer cell line drug response. Nature 560, 325–330 (2018).

8. A. Harms, E. Maisonneuve, K. Gerdes, Mechanisms of bacterial persistence during stress and antibiotic exposure. Science 354, (2016).

9. H. Nicoloff, K. Hjort, B. R. Levin, D. I. Andersson, The high prevalence of antibiotic heteroresistance in pathogenic bacteria is mainly caused by gene amplification. Nat Microbiol 4, 504–514 (2019).

10. V. Takhaveev, M. Heinemann, Metabolic heterogeneity in clonal microbial populations. Curr Opin Microbiol 45, 30–38 (2018).

11. E. A. Ottesen, J. W. Hong, S. R. Quake, J. R. Leadbetter, Microfluidic digital PCR enables multigene analysis of individual environmental bacteria. Science 314, 1464–1467 (2006).

12. A. Marusyk, V. Almendro, K. Polyak, Intra-tumour heterogeneity: a looking glass for cancer? Nat Rev Cancer 12, 323–334 (2012).

13. Y. S. Zhang, A. Khademhosseini, Advances in engineering hydrogels. Science 356, (2017).

14. Z. Zhu et al., Highly sensitive and quantitative detection of rare pathogens through agarose droplet microfluidic emulsion PCR at the single-cell level. Lab on a Chip 12, 3907–3913 (2012).

15. W. H. Tan, S. Takeuchi, Monodisperse alginate hydrogel microbeads for cell encapsulation. Advanced Materials 19, 2696-+ (2007).

16. P. Zimny, D. Juncker, W. Reisner, Hydrogel droplet single-cell processing: DNA purification, handling, release, and on-chip linearization. Biomicrofluidics 12, (2018).

17. C. J. Young, L. A. Poole-Warren, P. J. Martens, Combining submerged electrospray and UV photopolymerization for production of synthetic hydrogel microspheres for cell encapsulation. Biotechnol Bioeng 109, 1561–1570 (2012).

18. H. Ikehata, T. Ono, The Mechanisms of UV Mutagenesis. J Radiat Res 52, 115–125 (2011).

19. R. M. Wadowsky, S. Laus, T. Libert, S. J. States, G. D. Ehrlich, Inhibition of Pcr-Based Assay for Bordetella-Pertussis by Using Calcium Alginate Fiber and Aluminum Shaft Components of a Nasopharyngeal Swab. Journal of Clinical Microbiology 32, 1054–1057 (1994).

20. L. Xu, I. L. Brito, E. J. Alm, P. C. Blainey, Virtual microfluidics for digital quantification and single-cell sequencing. Nat Methods 13, 759–762 (2016).

21. X. Huang et al., Smartphone-Based in-Gel Loop-Mediated Isothermal Amplification (gLAMP) System Enables Rapid Coliphage MS2 Quantification in Environmental Waters. Environ Sci Technol 52, 6399–6407 (2018).

22. F. Schuler et al., Centrifugal step emulsification applied for absolute quantification of nucleic acids by digital droplet RPA. Lab on a Chip 15, 2759–2766 (2015).

23. Y. C. Tan, V. Cristini, A. P. Lee, Monodispersed microfluidic droplet generation by shear focusing microfluidic device. Sensor Actuat B-Chem 114, 350–356 (2006).

24. S. Haeberle et al., Alginate bead fabrication and encapsulation of living cells under centrifugally induced artificial gravity conditions. J Microencapsul 25, 267–274 (2008).

25. J. E. Kreutz et al., Theoretical design and analysis of multivolume digital assays with wide dynamic range validated experimentally with microfluidic digital PCR. Anal Chem 83, 8158–8168 (2011).

26. G. L. Francis, Albumin and mammalian cell culture: implications for biotechnology applications. Cytotechnology 62, 1–16 (2010).

27. R. D. Mitra, G. M. Church, In situ localized amplification and contact replication of many individual DNA molecules. Nucleic Acids Res 27, (1999).

28. S. J. Spencer et al., Massively parallel sequencing of single cells by epicPCR links functional genes with phylogenetic markers. Isme J 10, 427–436 (2016).

29. R. Zilionis et al., Single-cell barcoding and sequencing using droplet microfluidics. Nature Protocols 12, (2017).

30. L. M. Weber, C. G. Lopez, K. S. Anseth, Effects of PEG hydrogel crosslinking density on protein diffusion and encapsulated islet survival and function. J Biomed Mater Res A **90a**, 720–729 (2009).

31. Y. B. Wu, S. Joseph, N. R. Aluru, Effect of Cross-Linking on the Diffusion of Water, Ions, and Small Molecules in Hydrogels. J Phys Chem B 113, 3512–3520 (2009).

32. V. T. N. Tran et al., The sensitivity of real-time PCR amplification targeting invasive Salmonella serovars in biological specimens. Bmc Infectious Diseases 10, (2010).

33. T. Notomi et al., Loop-mediated isothermal amplification of DNA. Nucleic Acids Res 28, E63 (2000).

34. M. L. Xu, D. J. McCanna, J. G. Sivak, Use of the viability reagent PrestoBlue in comparison with alamarBlue and MTT to assess the viability of human corneal epithelial cells. J Pharmacol Tox Met 71, 1–7 (2015).

35. B. H. Neufeld, J. B. Tapia, A. Lutzke, M. M. Reynolds, Small Molecule Interferences in Resazurin and MTT-Based Metabolic Assays in the Absence of Cells. Anal Chem 90, 6867–6876 (2018).

36. J. Shemesh et al., Stationary nanoliter droplet array with a substrate of choice for single adherent/nonadherent cell incubation and analysis. Proc Natl Acad Sci U S A 111, 11293–11298 (2014).

37. E. Um, S. G. Lee, J. K. Park, Random breakup of microdroplets for single-cell encapsulation. Appl Phys Lett 97, (2010).

38. X. Lin, X. Huang, K. Urmann, X. Xie, M. R. Hoffmann, Digital Loop-Mediated Isothermal Amplification on a Commercial Membrane. ACS Sens 4, 242–249 (2019).

39. F. X. Fan, M. Y. Yan, P. C. Du, C. Chen, B. Kan, Rapid and Sensitive Salmonella Typhi Detection in Blood and Fecal Samples Using Reverse Transcription Loop-Mediated Isothermal Amplification. Foodborne Pathogens and Disease 12, 778–786 (2015).

40. L. B. Pinheiro et al., Evaluation of a Droplet Digital Polymerase Chain Reaction Format for DNA Copy Number Quantification. Analytical Chemistry 84, 1003–1011 (2012).

