## Supplementary Information for "A hydrogel beads based platform for single-cell phenotypic analysis and digital molecular detection"

**This PDF file includes:**

Supplementary Tables S1 to S2

Supplementary Figures S1 to S5

Supplementary Notes S1 to S3

### Supplementary Tables

| | | LAMP mix | PCR mix | TSB media |
|---|---|---|---|---|
| pH | | 8.8 | 8.3 | 7.3 |
| 7.5 w/v% | Sol-Gel transition time [min] | 7.0 | 11.5 | 43.0 |
| | | 8.5 | 13.5 | 53.0 |
| | Pore size | 27nm | | |
| 10 w/v% | Sol-Gel transition time [min] | 4.5 | 8.5 | 18.0 |
| | | 5.5 | 10.0 | 23.5 |
| | Pore size | 25nm | | |

**Table S1.** PEG hydrogel crosslinking characterization in bulk for LAMP mix, PCR mix, and TSB media. Sol-Gel transition time was experimentally determined (See **Materials and Methods**). The pH values were supplied by the manufactures. The pore sizes were estimated theoretically *(20)*.

| PCR | |
|---|---|
| Forward primer | 5’ CGCGAAGTCAGAGTCGACATAG 3’ |
| Reverse primer | 5’ AAGACCTCAACGCCGATCAC 3’ |
| Probe | 5’ FAM AAGACCTCAACGCCGATCAC 3’ |

| LAMP | |
|---|---|
| FIP | 5’ AACTTGCTGCTGAAGAGTTGGACCGAATGACTCGACCATC 3’ |
| BIP | 5’ CCTGGGGCCAAATGGCATTATGCACTAAGTAAGGCTGG 3’ |
| F3 | 5’ GACTTGCCTTTAAAAGATACCA 3’ |
| B3 | 5’ AGAGTGCGTTTGAACACTT 3’ |
| LF | 5’ TCGGATGGCTTCGTTCCT 3’ |
| LB | 5’ CAAGGGTTTCAAGACTAAGTGGTTC 3’ |

**Table S2.** Sequences of primers and probe for PCR and LAMP assays. PCR primers and probes target a region in gene STY0201 specific for *S.* Typhi for an amplicon size of 131 bp (*32*). LAMP primers target a 196 bp region within the *S.* Typhi specific gene STY1607 (*39*). The target regions for PCR and LAMP were both found to occur at one copy per cell, by searching the sequences within the complete genome of *S.* Typhi strain CT18 (Accession no. NC_003198) using Basic Local Alignment Search Tool (BLAST).

### Supplementary Figures

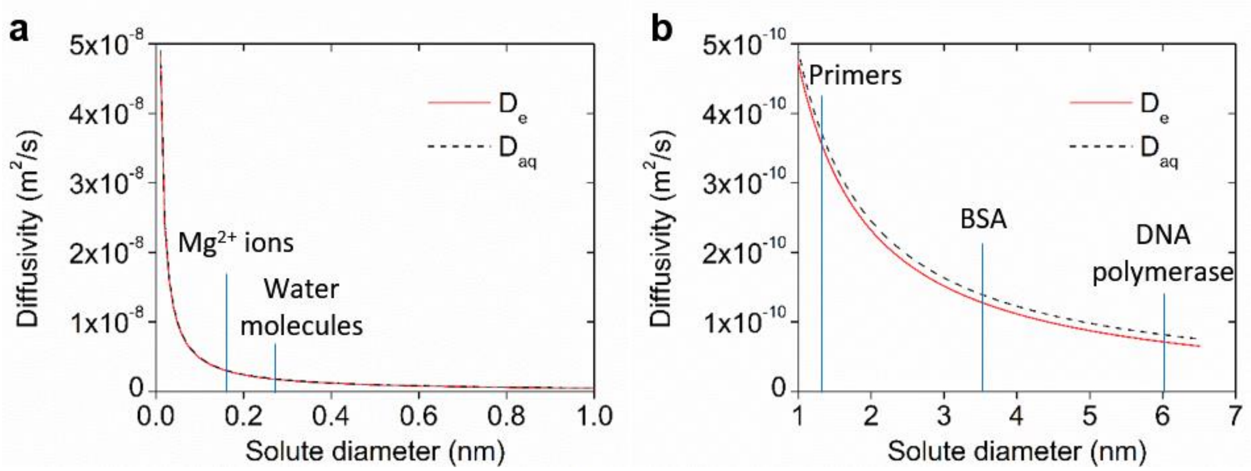


**Figure S1**. Calculated effective diffusivity ($D_e$) of solute in 7.5 w/v% PEG hydrogel matrix based on Weber et al. (*30*) and diffusivity in aqueous phase ($D_{aq}$) at 37 °C based on Stokes-Einstein equation as a function of solute hydrodynamic radius (**a**) from 0-1 nm and (**b**) from 1-7 nm. For functional molecules in the molecular assays approximate sizes are indicated on the plots.

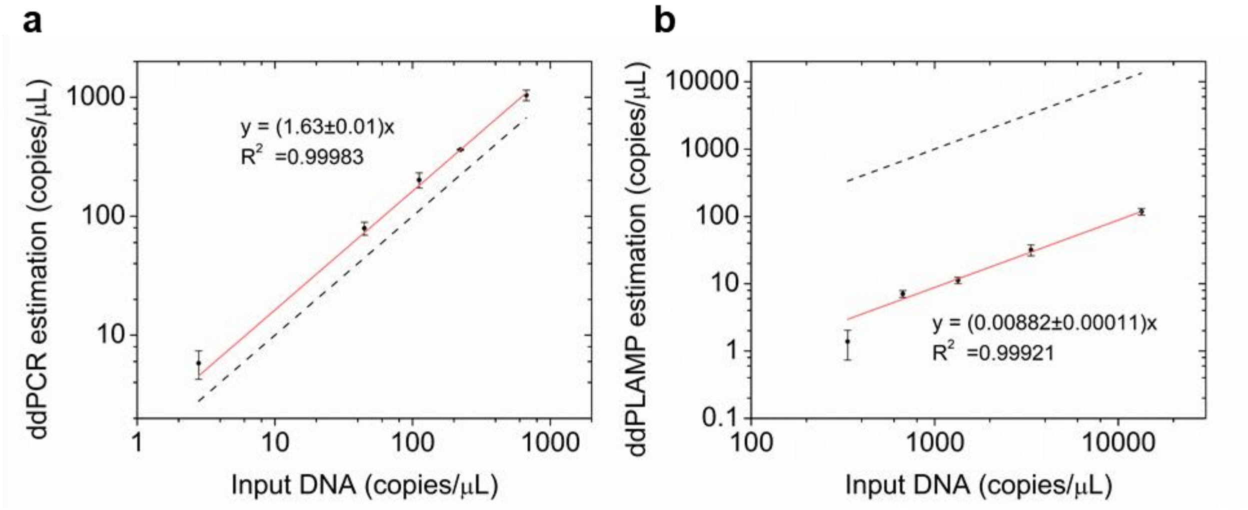


**Figure S2**. Estimated DNA concentration by (**a**) ddPCR for 24000, 1500, 600, 300, 100 times dilution of harvested S. Typhi DNA. and (**b**) ddLAMP for 200, 100, 50, 20, 5 times dilution of harvested *S*. Typhi DNA. compared with input DNA concentration. Input DNA concentration was calculated from dilution factor and OD600 measurement of cultured cells before DNA extraction using commercial kit. The dashed lines reference an exact match with input DNA concentration.

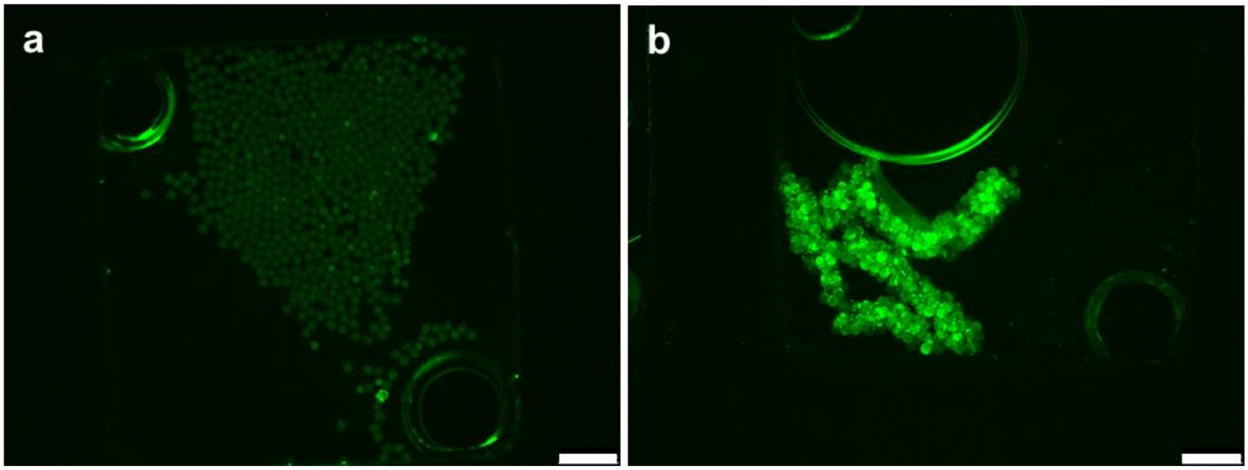


**Figure S3**. Example fluorescent images from preliminary gdLAMP experiments with (**a**) no target template and (**b**) 100 times dilution of harvested *S*. Typhi DNA. Scale bars, 1 mm. For preliminary gdLAMP experiments, LAMP MasterMix was used (New England BioLabs) instead of the customizable LAMP recipe specified in **Online Methods**. Each 20 μL of reaction mix for contained 1× LAMP MasterMix, 1.6 μM FIB and BIP, 0.2 μM F3 and B3, 0.8 μM LF and LB, 1× LAMP dye. 7.5 w/v% PEG hydrogel was added as 10× PEG monomers. The heating protocol involved 65 °C for 30 min and then 80 °C for 5 min. Aggregation of Gelbeads observed for positive samples but not for no-template controls. The extent of aggregation indicates occurrence of severe crosstalking.

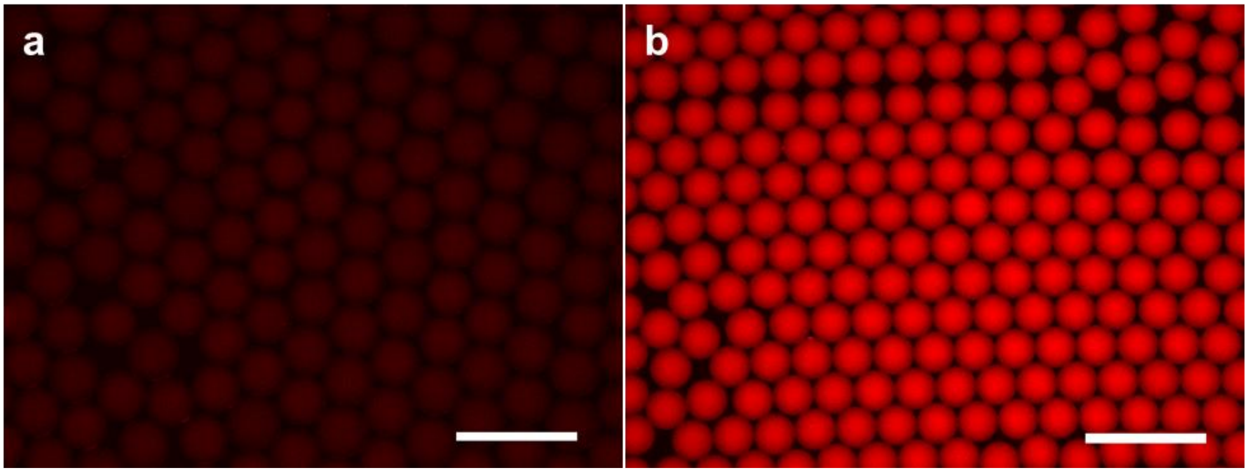


**Figure S4**. Example images of preliminary cell phenotyping experiments. With alamarBlue and media only, Gelbeads appeared to be much brighter than droplets before incubation even when imaged under a lower fluorescence (300 ms exposure for droplets and 25 ms exposure for Gelbeads). Scale bars, 500 µm.

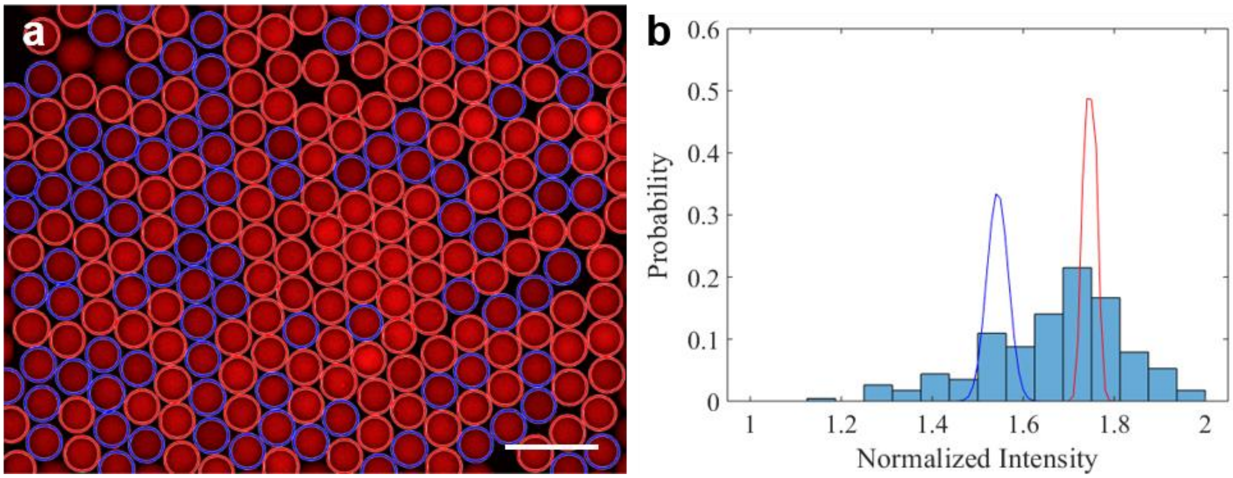


**Figure S5**. MATLAB analysis and threshold setting for images from phenotyping experiments. (a) An example image of phenotyping assay analyzed in MATLAB. Scale bar, 500 μm. The blue circles represent identified dark Gelbeads, and the red circles represent identified bright Gelbeads. (b) The histogram presents the occurrence probability of mean normalized intensity of Gelbeads analyzed based on the source image of (a). Gaussian fitting of the occurrence probability data generated two peaks, represented by the blue and red curves. The threshold suggested by this MATLAB script was set as the average of the mean (μ) of two peaks. This threshold was used to categorize negative and positive Gelbeads and produced the identification results on the left. We note that for 2 hours and 5 hours of incubation, the differences in normalized intensity of Gelbeads were too small for this thresholding method. In these cases, threshold enumeration and visual inspection were used instead for an approximately appropriate threshold.

### Supplementary Notes

#### Supplementary Note 1: Characterization of PEG hydrogel crosslinking

To identify the reasonable time frame for Gelbead generation, PEG gelation time in our targeted reaction matrices by bulk phase sol-gel transition experiments (online methods) was first measured. Ideally the compartmentalization process should be completed before the sol-gel transition starting, after which further crosslinking would considerably alter fluid properties such as viscosity and surface tension. For the three types of reaction matrix examined, results showed that the sol-gel transition start time spanned from 4.5 min to 43.0 min, and the crosslinking was accelerated by higher pH and higher monomer concentration (**Table S1**). Accordingly, the gelation time might be further extended by decreasing the crosslinking temperature. We then estimated if the lower gel concentration considerably affects the hydrogel properties. The theoretical pore sizes for our crosslinked PEG were close for 7.5% and 10% gel (**Table S1**), and both would allow diffusion of functional molecules in our applications including water, ions, small DNA fragments, and proteins that size from below an angstrom to ~ 6 nm (*31*). This estimation neglects non-ideality such as dangling ends or monomer self-interlinking, which might result in larger actual pore size. Using the same PEG monomers, 10 w/v% hydrogel concentration has been utilized in cell encapsulation, multiple displacement amplification (MDA) and LAMP (*20, 21*). We reason that lowering the hydrogel concentration to 7.5 w/v% would benefit our applications by allowing more time for Gelbead generation and creating looser hydrogel network for reagent diffusion. Therefore, 7.5 w/v% PEG was used in further experiments.

**Supplementary Note 2**: **Droplet generation performance and sources of error**

To achieve high throughput analysis of droplets with limited available instruments, we chose to analyze droplets through fluorescence imaging. As in the reported protocol, droplets were extracted into a viewing chamber, made by bonding a commercial plastic chamber onto a glass slide, for fluorescence imaging at 1.25× objective. We acknowledge that this protocol might have introduced systematic error in size characterizations by 1) pipetting droplets from the microcentrifuge tube into the viewing chamber, 2) noise difference from focus point to the edges within an image, and 3) image processing bias by MATLAB when identifying the circular-shaped edges. These sources of error would lead to overestimation of size distribution. Therefore, it is anticipated that the actual CV of the generated droplets or Gelbeads should be lower than reported. Generally, the reported sizes and CVs of generated droplets, optimized at 175 µm diameter with CV of 5%, are comparable to those generated by centrifugal microfluidics reported in literature. For example, Haeberle et al. used polymer-tube micronozzles for alginate bead generation, and reported beads generated at diameters tunable from 180 to 800 µm with CV of 7–16% (*24*). As another example, using a lab-on-a-disk centrifugal droplet generation with, Schuler et al. reported droplet diameters of 120 to 170 µm with CV of 2–4%. It should be noted that these values represent only 20 measurements of droplet under high microscope objective (*22*).

**Supplementary Note 3: Microscope objective choice**

For ddPCR and gdPCR, 5× objective was used in fluorescent microscope imaging to more accurately capture the assay quantification performance. Taqman probe, commonly used in PCR for enhanced detection specificity by hydrolysis upon encounter of specific sequence target, was employed in our PCR assay recipe. In this case, the fluorescence exhibited by negative compartments was too low to be distinguishable from the oil phase under 1.25× objective. To be able to count negative compartments, a zoomed-in view using 5× objective had to be used instead of 1.25× objective, which was employed to image ddLAMP and gdLAMP results. A commercial DNA-intercalating dye was used in LAMP so that the higher background fluorescence was observed in negative compartments. 1.25× objective could image an area of 0.785 $cm^2$ to include the whole viewing chamber at one image shot. The smaller view at 5× objective was compensated by taking 5 images of different areas in the viewing chamber. A similar strategy was applied in Gelbeads imaging for phenotyping experiments to better distinguish the Gelbeads with varying fluorescence levels.
